# Evaluation of deep and dynamic proteomic screening strategies at sub-50Hz scan rate and without automation

**DOI:** 10.64898/2026.04.15.718630

**Authors:** Bhavesh S. Parmar, Yuanyuan Liu, Parviz Ghezellou, Christian Münch

**Affiliations:** Institute of Molecular Systems Medicine, Goethe University, Frankfurt am Main 60528 Germany

## Abstract

Advances in ultra-fast mass analyzer technology and procedural automation have enabled proteomics screening at the throughput of hundreds of proteomes per day. However, these approaches often require expensive instrumentation upgrades and robotic automation that remain inaccessible to many research laboratories and core facilities. In this study we address the feasibility of scaling up proteomic screening capabilities with minimal upgrade cost by focusing on (a) strategies for non-automated high-throughput sample preparation from 96-well cell culture, (b) data acquisition on sub-50Hz scan speed hybrid and tribrid Orbitrap instruments and (c) data analysis strategies for label-free and labeled proteomic screening. We find that the 96-well format STrap, in combination with C18 plates, provides the most robust throughput for a non-automated sample preparation workflow. Furthermore, we show that for static proteomes, an isobaric tandem mass tag-based (TMT) multiplexing approach provides deeper and more precise proteome coverage whereas label-free data-independent acquisition (DIA) is more accurate, albeit with a reduced dynamic range and more missing values. Finally, we extend the optimized workflow to proteome turnover studies using pulsed stable isotope labeling by amino acids in cell culture (pSILAC), highlighting the key advantages and trade-offs of DIA and TMT data-dependent acquisition strategies for capturing protein translation. Together, these results provide a practical framework for designing high-throughput proteomics experiments that balance throughput, depth, and quantitative accuracy using existing instrumentation, without requiring major hardware upgrades or automation.

## Introduction

Over the past two decades, mass spectrometry (MS)-based proteomics has matured into a cornerstone technology for biological and biomedical research^1–3^. This progression has been multifaceted, driven not only by advances in instrumentation and computational analysis, but also by the expansion of training programs, shared core facilities, and standardized workflows that have broadened accessibility across academic, clinical, and industrial sectors. Routine bottom-up proteomics is now widely adopted, supported by robust and reproducible protocols and end-to-end commercial kits that lower experimental barriers and improve inter-laboratory reproducibility^4^. Advanced liquid handling robots have further enabled automated high-throughput sample processing within relatively short time frames^5–9^. On the acquisition side, recent innovations-including ion mobility separation, mass analyzers operating at scan rates exceeding 200 Hz^10,11^, and robust liquid chromatography systems-have enabled analytical capacities approaching 500 samples per day^12^. Notably, deep proteome coverage from complex biological matrices can now be attained with sample runtime as brief as 15 minutes and 10$ of user cost^13,14^. Consequently, fully unmanned proteomics workflows encompassing sample preparation and data acquisition have become technically feasible^15^. Despite these advances, the practical accessibility of cutting-edge proteomics remains uneven. While the field increasingly favors time-of-flight-based instrumentation and greater end-to-end automation, a substantial proportion of laboratories-particularly those operating under resource limitations as of 2024— continue to rely on Quadrupole-Orbitrap (Q-OT) and Quadrupole-Orbitrap-Linear Ion Trap (Q-OT-LIT) instruments with moderate scan rates (<50 Hz)^16^ and non-automated methods.

In this study, we systematically evaluate sample preparation strategies, data acquisition modes and data-analysis approaches for both labeled and label-free proteomics, with the explicit goal of maximizing throughput and quantitative depth using Orbitrap-based instruments operating below 50Hz scan rates and without reliance on automation. Our central objective is to define realistic, accessible proteomics workflows that can be implemented within standard cell and molecular biology laboratories, enabling high-throughput proteome measurements akin to cell screening platforms.

Specifically, we first assess high-throughput sample preparation strategies compatible with 96-well cell culture formats, comparing in-solution (adapted from Simplit^17^ i.e. Simplit2), filter-based (STrap^18^), and on bead aggregation (SP3^19^) based enzymatic digestion. We then benchmark label-free quantitative data-independent acquisition (LFQ-DIA) and tandem mass tag–based data-dependent acquisition (TMT-DDA) -across multiple Orbitrap platforms, including the Q-Exactive HF, Orbitrap Fusion Lumos Tribrid and Orbitrap Ascend Tribrid, evaluating achievable depth, throughput, and quantitative accuracy -within a 24-hour experimental window. Lastly, we integrate 96-well cell culture with pulsed stable isotope labeling by amino acids in cell culture (pSILAC;2SILAC and 3SILAC formats) to assess DIA-and TMT-based approaches for capturing temporal proteome dynamics in medium- to high-throughput fashion. Collectively, this work provides a practical and adaptable framework for performing deep and dynamic proteomic screens using widely available Orbitrap instrumentation and non-automated workflows. The strategies outlined here are readily transferable to diverse biological systems-including adherent cells, suspensions cultures or tissues, and can be applied across multiple research contexts, such as clinical proteomics, chemoproteomics, and CRISPR- or siRNA-based functional proteomics screens. By aligning methodological rigor with infrastructural realities, this study aims to bridge the gap between state-of-art proteomics capabilities and their widespread, routine implementation.

## Materials and methods

### Cell culture and SILAC labelling

HeLa cells were used for all the experiments and grown in RPMI 1640 or SILAC RPMI 1640 (Gibco™ - Product number 11875093 and 15477983) supplemented with 10% fetal bovine serum and appropriate amino acid isotopes (Silantes) (for SILAC). For sample preparation optimization, 20,000 HeLa cells/well were seeded in 96-well cell culture plate (Sarstedt) and grown overnight in 100µL of medium. The medium was decanted and adhered cells gently washed 3X with 200µL Phosphate buffer saline (PBS) and frozen with 75µL of lysis buffer (see below). For dynamic proteome with Thapsigargin/DMSO treatment, 20,000 cells/well were seeded in a 96-well plate and grown overnight in 100µL of SILAC Light (K0,R0) medium. Following that, the light medium was decanted, cells rinsed with 200µL PBS once and treated either 1µM Thapsigargin or Dimethyl Sulfoxide (DMSO) in Heavy (K8, R10) medium for 2 hrs. The cells were then washed 3X with PBS and one batch (plate) was frozen for 2SILAC experiment in 75 µL of lysis buffer. Another plate was further incubated with SILAC intermediate (K4, R6) medium for 2 hrs, rinsed 3X with PBS and frozen in 75µL of lysis buffer for 3SILAC experiment.

### High-throughput sample preparation from 96-well cell cultures

For STrap and SP3, lysis buffer contained 5% sodium dodecyl sulfate (SDS), 5mM tris(2-carboxyethyl)phosphine (TCEP), 25mM chloroacetamide (CAA) and 1X protease inhibitor cocktail (Roche cOmplete EDTA free - 11836170001) in 100mM triethylammonium bicarbonate (TEAB). For Simplit2, lysis buffer contained 2% sodium deoxycholate (SDC), 5mM TCEP, 25mM CAA and 1x protease inhibitor cocktail in 100mM TEAB. For all samples described in Fig.1 and Fig.3, plates were thawed by snap heating at 95°C for 10 mins in Thermomixer C (Eppendorf) and spun at 1500xg. Following that, the samples were sonicated using 8-horn probe sonicator (Qsonica part no.4599) connected to ultrasonic processor (Sonics vibra-cell VCX130) at 40% amplitude for 2 mins with 1s on 1s off pulse. The samples were reduced with 1µL of 500mM TCEP and alkylated with 5µl of 500mM CAA, both for 20mins at RT. For Simplit2 protocol, 0.2µg of trypsin+lys-C mix (1:1) (Promega V5111 and VA1170) was added directly to the plate, sealed with a silicon mat between the plate and the lid, and incubated overnight at 37°C. The digested peptides were acidified with 200µL of 99% isopropanol/ 1% trifluoroacetic acid and loaded onto Styrenedivinylbenzene-reverse phase sulfonated (SDB-RPS) plate (Affinisep-AttractSPE microelution μ96W-RPS-M.T.1.1) attached to 1.2 mL deep well plate and spun at 600 x g for 2 mins. The peptides were washed once with 200 µL of 99% isopropanol/1% trifluoroacetic acid, and 200µL of 2% acetonitrile/0.1% trifluoroacetic acid by spinning at 1500xg for 1 min. The cleaned peptides were eluted in 80% acetonitrile/5% ammonia and dried in a vacuum concentrator. For STrap plate, the samples were acidified with 5 µL 25% phosphoric acid, and the lysate was transferred into 1.2 mL deepwell plate (Abgene™ AB1127). 700 µL of 90% methanol, 100 mM TEAB was added to the lysate and loaded onto STrap (ProtiFi - 96 well micro plate) in 2 batches of 390 µL and spun at 1500g for 1 min with a 1.2mL deepwell plate underneath. The samples were washed 3X with 200 µL 90% methanol, 100 mM TEAB (Sigma-Aldrich T7408) by centrifugation at 1500 x g for 1 min each. Following that, 0.2 µg of trypsin+lys-C mix (1:1) in 125 µL of 100 mM TEAB was added to the filter and incubated overnight at 37°C on a fresh 1.2mL deep well plate. The digested peptides were eluted from STrap along with 80µL of 100mM TEAB first, then with 80µL of 0.2% trifluoroacetic acid and lastly with 80µLof 50% acetonitrile/ 0.2% trifluoroacetic acid. The peptides were dried in a SpeedVac before being resuspended in 100 µL of 0.2% trifluoroacetic acid for C18 clean up. For SP3, the lysate was transferred to a 1ml LoBind plate (Eppendorf - 0030504208) and 5 µL of 50 µg/µL prewashed beads (1:1 Cytiva SpeedBead Magnetic Carboxylate - 6515,4515) were added, followed by 100 µL of ethanol and vortex for 5mins at 1200 rpm. The plate was set on a magnetic rack (LifeSep Z740158) for 10 mins to allow the beads to separate and the liquid was decanted using a multi-channel pipette. This process was then repeated 3X with 100 µL of 80% acetonitrile. The beads with protein aggregates were then resuspended in 100µL of 0.2µg trypsin+lys-c mix, briefly sonicated to detach the beads and incubated on Thermomixer at 37°C overnight with intermittent shaking at 1200rpm for 2 min every 1 hr. The digested SP3 peptides were acidified to 0.2% trifluoroacetic acid and spun at 1500xg for 2 mins before setting on a magnetic rack for 10mins and collecting the acidified peptides for C18 clean up. All C18 plate (Affinisep-AttractSPE microelution μ96W-C18-M.T.1.1) spins were performed at 600xg for 3 min with 1 ml deep well plate attached underneath. First, the C18 was washed with 200µL 50% acetonitrile/0.2% trifluoroacetic acid and equilibrated 3X with 200µL of 0,2% trifluoroacetic acid. The STrap and SP3 samples were loaded and washed 3X with 200µL 0.2% trifluoroacetic acid followed by elution with 50% acetonitrile/ 0.2% trifluoroacetic acid and drying in vacuum concentrator. The samples were then resuspended in either 100mM TEAB for the microBCA peptide assay or 2% acetonitrile/ 0.1% trifluoroacetic acid for LC-MS/MS analysis.

**Fig 1:**
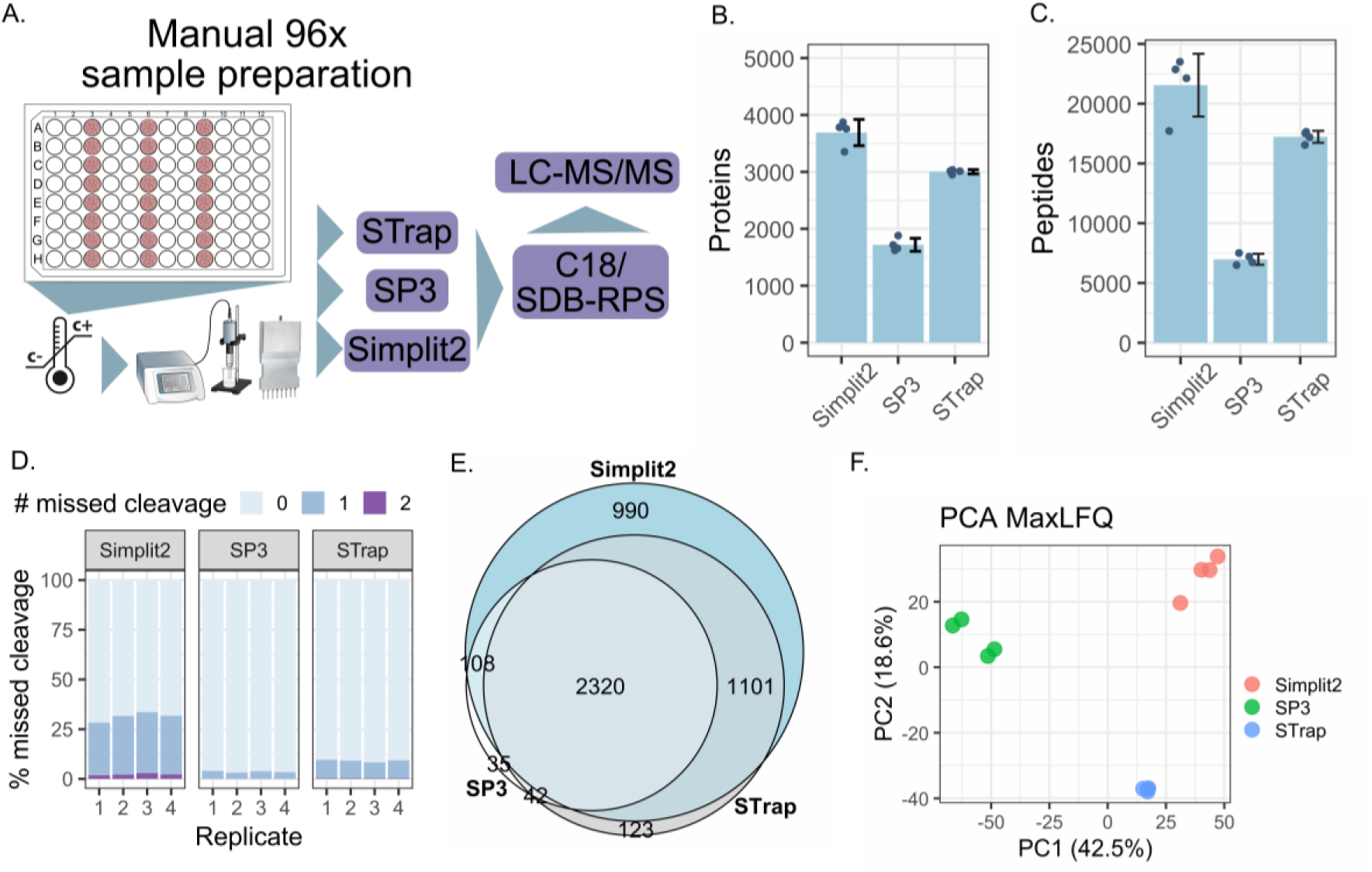
Comparison of high-throughput, non-automated sample preparation methods for cells grown in 96-well plate. A. Experimental scheme of cell culture and sample preparation methods compared for manual processing. 20,000 cells were seeded and grown overnight, followed by freeze-thaw lysis, homogenization using multi-probe sonication and digestion using STrap, SP3 or Simplit2 protocol followed by C18 or SDB:RPS cleanup B. Proteins and C. Peptides identified across across replicates from 125ng of digest using DDA-LFQ method. D. Proportion of trpytic missed cleavages across the three methods. E. Overlap of proteins identified across the three methods. F. Principal component analysis of MaxLFQ intensities per replicate showing inter sample variance within each compared method.

### Three-species proteome benchmarking mixtures

Commercially purchased peptide digests for bacteria (*Escherichia coli*, MassPREP-Waters 186003196), yeast (*Saccharomyces cerevisiae*, Promega V7461) and human (K562, Promega V6951) were resuspended in 10% acetonitrile to a concentration of 2ug/µL and pooled at different human:bacteria:yeast ratios (ctrl - 65:15:20, Mix1 - 65:30:5, Mix2 - 65:20:15) in triplicate. For LFQ, the pooled digests were diluted to 1% acetonitrile/0.2% trifluoroacetic acid and stage-tipped (EMPORE 3M 2215-C18), same as C18 plates, but with centrifugation at 1500xg. For TMT labelling, samples were diluted to achieve 100mM 3-[4-(2-hydroxyethyl)piperazin-1-yl]propane-1-sulfonic acid (EPPS) in 10% acetonitrile, labelled with TMT10-plex, cleaned up using C18 Sep-Pak cartridge (tC18, 50 mg, Waters WAT054960) and fractionated by high-pH C18 reversed-phase chromatography (see below).

### pSILAC experimental design for dynamic proteome measurements

The dynamic SILAC plates were processed in the same fashion following the above-mentioned STrap protocol followed by C18 clean up in 96-well format. For DIA analyses, samples were resuspended in 15µL of 2% acetonitrile, 0.1% trifluoroacetic acid and 3µL was injected. For TMT were resuspended in 20µL of 100mM EPPS, 10% acetonitrile, cleaned up, fractionated and run on Ascend and Lumos.

### Isobaric labelling with TMT and TMTpro

For 3 species mix, 10 µg per sample was labelled with TMT10 in the following order of replicates per condition-Ctrl (127N,127C,128N), Mix1 (128C,129N,129C), and Mix2 (130N,130C,131N). For dynamic 2SILAC and 3SILAC combined with TMT, the samples were labelled with TMTpro such that all replicates per condition get the same tag (N or C) – DMSO (127N-134N), Thapsigargin (126C-133C). Dried 5 mg TMT stock was resuspended in 250 µL of 100% acetonitrile and 1 µL was added to the above-mentioned samples and incubated for 1 hr at 24 °C. Following that, the reaction was quenched with 0.5% hydroxylamine (final concentration) for 15 mins at 24°C. 1 µL of each TMT channel was pooled to confirm the labelling efficiency via LC-MS/MS. Following confirmation of >99% labelling, the channel volumes were calculated based on median S/N and pooled together.

### High-pH reversed-phase peptide fractionation

The pooled TMT channels were acidified to 0.2% trifluoroacetic acid and cleaned with C18 SepPak cartridge (tC18, 50 mg, Waters) with 3X 2% or 5% acetonitrile/ 0.1 % trifluoroacetic acid (for TMT11 and TMTpro, respectively), eluted with 60% acetonitrile and dried in a vacuum concentrator. The dried TMT11 and TMTpro samples were resuspended in 40 µl 5 mM ammonium bicarbonate/ 5% acetonitrile, sonicated in a water bath for 1 min and concentration measured using microBCA. Sample concentration was adjusted to 1.5 µg/µL and 45 µg of the sample was used for high-pH reversed-phase fractionation on an Ultimate 3000 HPLC system (Thermo Fisher Scientific) equipped with a C18 column (XSelect CSH, 1 mm × 150 mm, 3.5 µm particle size; Waters SKU186005251). Peptides were separated using a multistep gradient from 3-60% Solvent B (5 mM ammonium bicarbonate in 80% acetonitrile) over 65 min at a flow rate of 30 µL/min. Eluting peptides were collected every 43 seconds from minute 2 for 69 minutes into a total of 96 fractions, which were concatenated into 24 or 12 fractions. Pooled fractions were dried in a vacuum concentrator and resuspended in 2% acetonitrile/ 0.1% trifluoroacetic acid for LC-MS analysis.

### LC-MS/MS data acquisition

Detailed acquisition parameters for all DDA and DIA methods are provided in Supplementary Table S1 (Sheets 1 and 2). All LC–MS analyses were performed using nanoLC systems (Easy-nLC 1200 or Vanquish Neo; Thermo Fisher Scientific) operated at flow rate of 400 nL/min and coupled to hybrid Orbitrap mass spectrometers (Q-Exactive HF, Orbitrap Fusion Lumos, and Orbitrap Ascend). Mobile phases consisted of Solvent A (0.1% formic Acid in water) and Solvent B (80% acetonitrile with 0.1% Formic Acid). Samples were loaded at 950 bar pressure using 6µl of Solvent A onto an in-house packed 35 cm fused-silica column (75µm inner diameter) packed with 1.9µm Reprosil-Pur C18 particles (Dr. Maisch r119.b9.). For experiments shown in Fig. 1, data acquisition was performed on a Q-exactive HF coupled to an Easy-nLC 1200. Approximately 125 ng of peptides (quantified using a microBCA assay) were injected and separated over a 90 min gradient consisting of 4-20% B in 60 min, 30-40% B in 20 min, 40-60% B in 5 min followed by 9 mins of column wash at 95% B. For three-species mixture experiments using TMT-MS3 acquisition on Easy-nLC 1200 coupled to Orbitrap Fusion Lumos, approximately 500 ng per fraction was loaded and separated using a 90 mins gradient (5-10% B in 3 min, 10-28% B in 72 min, 28-40% B in 15 min, 40-60% B in 5 min followed by 9 mins of column wash at 95% B). For three-species mixture experiments with TMT-MS2 and TMT-MS3 [real-time search (RTS) and real-time search closeout (RTScl)] on the Vanquish Neo coupled to Orbitrap Ascend, approximately 250 ng per fraction was injected and separated over 90 mins gradient (7-28% B in 70 min, 28-40% B in 13 min, 40-60% B in 5 min followed by 6 min of column wash at 95% B). For real-time search (RTS) experiments, SPS-MS3 triggering was enabled using a custom three-species FASTA database containing one gene per protein sequence for each organism downloaded from UniProt. RTS searches were performed either with (RTScl) or without (RTS) peptide/protein closeout, with the closeout threshold set to two peptides per protein. RTS parameters included: TMT on peptide N-terminus and lysine as fixed modifications and Methionine oxidation as variable modification, XCorr cut-off of 1 and deltaCn 0.1 at 5 ppm. For DIA experiments, 250 ng (Orbitrap Ascend) and 500 ng (Q-Exactive HF) of peptides were injected and separated using 45, 60, 90 or 120 min gradients with optimized isolation window schemes and instrument-specific acquisition parameters (Supplementary Table S1). For DIA with pSILAC labelled samples (2SILAC and 3SILAC, n = 4 biological replicates per condition), 500 ng was loaded on the column, and the same 120-minute gradient was used on Q-Exactive HF and Lumos (both online with Easy-nLC 1200) with MS1 and DIA scans were acquired at 120,000 and 30,000 resolution (at *m/z* 200), respectively (Supplementary Table S1). For pSILAC-TMTpro experiments, data were acquired using MS3 on the Orbitrap Fusion Lumos and MS3-RTS on the Orbitrap Ascend (n = 8 biological replicates per condition). Identical acquisition settings were used on both instruments, with the Targeted Mass Difference (TMD) node enabled to detect either two or three isotopic states (2SILAC and 3SILAC, respectively), with the key difference being MS3 resolution (45K or 50K) and maxIT (91ms or 86ms), respectively, with 250ng of sample loaded on the column. The fractions were separated over a 120 min gradient (7-25% Solvent B ramp in 90 min, followed by 40% in 30 min and 60% Solvent B in 5 min). RTS for pSILAC-TMTpro experiments was set by adding heavy and intermediate lysine and arginine SILAC masses as variable modifications, together with Methionine oxidation, two variable modifications allowed per peptide, and a human UniProt FASTA database (one gene per sequence). All other RTS parameters were identical to those used for the three-species experiments.

### Proteomic data processing and statistical analysis

DDA data were analysed using Proteome Discoverer (v2.4 or v3.2) or FragPipe v23.0, while DIA data were processed using DIA-NN v2.2. Across all search engines, Cysteine carbamidomethylation (+57.02146 Da) was specified as a fixed modification, while protein N-terminal acetylation (+42.0106 Da) and Methionine oxidation (+15.9949 Da) were included as variable modifications. For TMT experiments, TMT labeling on peptide N-terminal and lysine residues was specified using mass shift of +304.20715 Da (TMTpro) and +229.16293 Da (TMT classic). For the sample preparation comparison experiments, “Basic search” workflow in FragPipe was used with default parameters and match-between-run (MBR) disabled. All TMT experiments were searched on FragPipe with default settings for MS2 or MS3 reporter ion quantification and searched against either a concatenated three-species FASTA database or Human UniProt FASTA database (one gene per sequence). For pSILAC-TMTpro experiments, the default SILAC workflows in FragPipe were adapted to include TMT and SILAC+TMT masses as variable modifications, and “Run Isobaric Quant” was enabled in the Quant (Isobaric) tab. For both three-species and SILAC DIA search on DIA-NN, first a spectral library was generated with default parameters using either the three-species concatenated FASTA or Human UniProt FASTA database and then raw files were subsequently searched against the resulting predicted spectral library (predicted.speclib file). For pSILAC DIA analysis, additionally, --fixed-mod SILAC,0.0,KR,label --lib-fixed-mod SILAC --channels SILAC,L,KR,0:0; SILAC,H,KR,8.014199:10.008269 --original-mods --channel-run-norm for 2SILAC and --channels SILAC,L,KR,0:0; SILAC,M,KR,4.025107:6.020129;SILAC,H,KR,8.014199:10.008269 --original-mods --channel-run-norm for 3SILAC was added into “Additional options”. The resulting report.parquet files were processed in RStudio using the DIA-NN R package. All DIA results were filtered for Q-value, PG.Q-value, Lib.Q-value, Lib.PG.Q-value and channel.Q-value (for SILAC), all set to < 0.01, along with Quantity.Quality > 0.8 before protein roll up. The number of data points per chromatographic peak was estimated by dividing the retention time width of precursor signals by the maximum cycle time for each acquisition method. For pSILAC DIA the precursors from report.parquet from DIA-NN were separated based of channel assignment and for pSILAC-TMT, PSMs were separated based on peptide modification. FragPipe PSM.txt outputs were filtered for Purity > 0.5 and Peptide probability > 0.9, and the lowest 5% of PSM intensities were removed prior to protein summarization using parsimony-based inference. PSM output from Proteome Discoverer was filtered for isolation interference (>0.5), SPS match percentage (>60) and PSM uniqueness before summing signal-to-noise ratios for Protein rollup. For all TMT results, the filtered PSM intensities were normalized to total reporter ion intensity per TMT channel before protein summarization, and for DIA PG.MaxLFQ intensities were used for downstream analysis. Differential expression was evaluated using Limma, gene ontology enrichment using ClusterProfiler, and visualization by ggplot2 in RStudio.

## Results

### Comparison of high-throughput and non-automated sample preparation methods for 96-well cell culture

We systematically evaluated three non-automated, high-throughput sample preparation workflows compatible with 96-well cell culture formats: Simplit2 (an adaptation of the Simplit protocol^17^), SP3^19^ and STrap^18^ (Fig.1A). These methods differ substantially in reagent cost, hands-on complexity, digestion efficiency and -time required for sample preparation as investigated in detail by Varnavides et al^4^. Cells in individual wells were lysed by freeze-thawing, sonicated with a multi-horn probe and digested with the aforementioned methods followed by clean up with C18 (for SP3 and STrap) or SDB-RPS stage tips (for Simplit2). Evaluating sample measurements across replicates, Simplit2 and STrap yielded higher numbers of protein and peptide identifications relative to manual SP3 processing (Fig.1B and C). Of importance, Simplit2 produced a higher proportion of missed cleavages as compared to SP3 and STrap (Fig.1D). Although STrap mini plates are typically recommended for protein input between 100-300µg (by the vendor), we evaluated their performance at lower inputs to assess suitability for 96-well cell culture experiments. Using HeLa lysate inputs ranging from 5-100µg, we observed consistent protein and peptide identification numbers across all inputs, with only a marginal increase in variability at 5µg (Fig.S1A-C). Moreover, STrap coupled with C18 clean up provided a higher peptide recovery when compared with Simplit2 combined with SDB-RPS or SP3 combined with C18 clean up (Fig.S1D). The identified proteins were consistent across the three methods (Fig.1E) with Simplit2 producing relatively larger peptides (Fig.S1E) owing to higher missed cleavage proportion (Fig.1D). Importantly, quantitative reproducibility across replicates was highest for STrap, which exhibited the lowest coefficient of variation relative to Simplit2 and SP3 (Fig.1F, Fig.S1F). We further assessed compatibility of STrap and Simlpit2 with isobaric labeling (as shown in the original Simplit publication^17^). While unfractionated TMTpro samples show ∼80% overlap in proteins measured with both methods (Fig.S1G), the results are in concordance with our LFQ observations with less inter-sample variation observed with STrap as compared with Simplit2 (Fig.S1H-I). Overall, these results demonstrate that both STrap and Simplit2 are well suitable for non-automated 96-well proteomic sample preparation workflows, whereas SP3 may require further optimization for manual 96-well plate handling, especially for <10µg inputs. Between STrap and Simplit2, the former offers better digestion efficiency and quantitative reproducibility, whereas the latter is relatively simpler and more economical for high throughput proteomic screens at the expense of increased inter-sample variability and a higher proportion of missed cleavages.

### Comparison of data acquisition strategies for deep global proteomics

To benchmark data acquisition strategies under constrained scan-rate conditions, we analyzed three-species peptide mixtures spanning a broad dynamic range of peptide abundances using either label-free data-independent acquisition (DIA) or tandem mass tag (TMT)–based data-dependent acquisition (DDA) workflows. Peptide digests derived from bacteria, yeast, and human proteome were combined in defined ratios (Fig. 2A) to generate control (ctrl), mix1 and mix2 sample mixes (*n* = 3 replicates/sample mix). For label-free DIA, each mixture was injected individually and analyzed using four different chromatographic gradient lengths on both the Orbitrap Ascend and Q-Exactive HF mass spectrometers, in order to estimate the sample throughput and proteome depth achievable for both instrumental setups. We selected fixed isolation-windows schemes that resolved the spectral complexity, preserved at least six median data points per chromatographic peak and avoid trapezoidal quantitation^20^ (Fig.S2A). The data point per precursor were estimated using precursor RT.Start and RT.Stop, the maximum attainable cycle time and median full-width at half-maximum (FWHM) values for each LC-MS configuration (Fig.S2B-C). For isobaric TMT-based DDA, samples were separated by high-pH reversed-phase fractionation and analyzed either on an Orbitrap Lumos using MS3 acquisition with 24 fractions (90-min gradients) or on an Orbitrap Ascend using 12 fractions (90-min gradients) with MS2, MS3 with real-time search (MS3RTS), or MS3RTS with closeout limiting MS3 sampling to two peptides per protein per fraction (MS3RTScl) (Fig. 2A). Detailed acquisition parameters and instrument settings are provided in the Materials and Methods section and Table S1.

**Fig 2:**
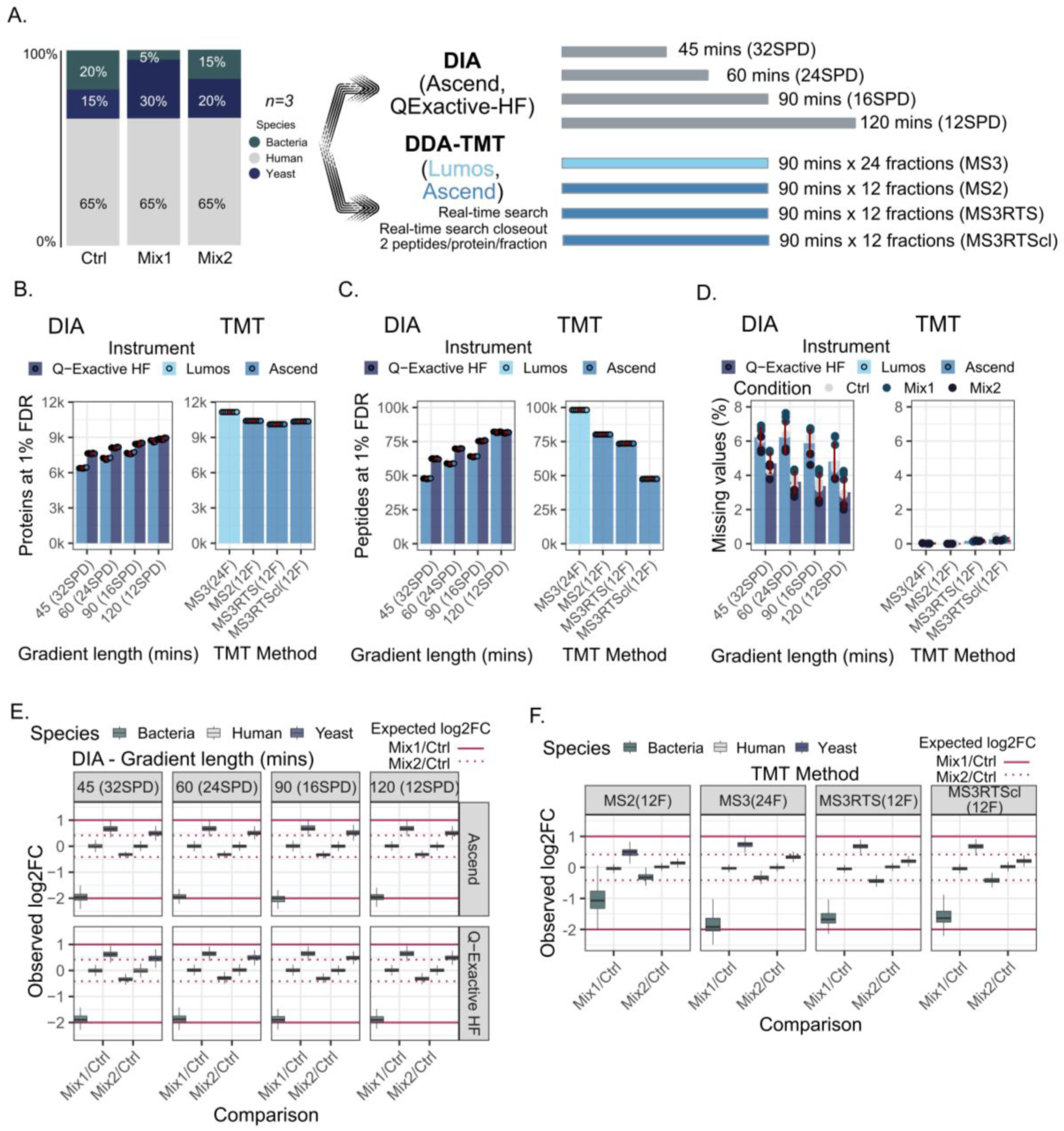
Comparison of data acquisition strategies for deep global proteomics A. Tryptic peptide digests of yeast and bacteria were mixed with human background in three different ratios (namely Ctrl, Mix1 and Mix2) to create a dynamic range of difference of 2-fold (yeast - Mix1 vs Ctrl), 4-fold (bacteria - Mix1 vs Ctrl) or 1.33-fold (yeast and bacteria - Mix2 vs Ctrl). The samples were run individually (n=3 technical replicates per sample) either as DIA on Q-Exactive HF and Ascend with 4 different gradient lengths amounting to 4 levels of throughput or labeled with TMT, fractionated into 24 or 12 fractions and run over a 90-mins gradient with SPS-MS3 (on Lumos - light blue) or MS2, Real-time search (RTS) triggered SPS-MS3 and RTS triggered SPS-MS3 with 2 peptides/protein/fraction close out option on Ascend (dark blue). B. Proteins and C. Peptides quantified at 1% FDR for all the compared DIA and TMT methods with dots indicating identifications per replicate. D. percentage of missing values per method with dots corresponding to sample type (Ctrl, Mix1, Mix2) and error bars showing standard deviation across all 9 samples. E. Boxplot showing distribution of log2 fold change for each comparison (Mix1/Ctrl and Mix2/Ctrl) across all methods compared with black line within each box indicating the median observed log_2_ fold change and red lines at expected log_2_ fold change (solid for Mix1/Ctrl and dotted for Mix2/Ctrl).

Under these conditions, fractionated TMT -DDA consistently yielded greater proteome depth than single-shot, library-free DIA. Ranked by depth, -MS3 with extensive fractionation (24 fractions) performed best, followed by MS2 (12 fractions), MS3RTScl (12 fractions) and MS3RTS (12 fractions) (Fig.2B-C). With equivalent acquisition time, the three TMT approaches using 12 × 90-min fractions quantified >10,000 proteins (mean peptide counts per TMT channel ≈ 80,180, 73,519 and 47,569 for MS2, MS3RTS and MS3RTScl, respectively), as compared to <9,000 proteins with 81,937 and 81,509 peptides for DIA on Ascend and Q-Exactive HF, respectively (Fig.2B-C). TMT-MS3 without RTS on the Lumos achieved the deepest coverage (>11,158 proteins and 98,171 peptides) but required twice the acquisition time. As expected, the quantified proteins and peptides progressively declined with shorter gradient lengths in DIA, with a more pronounced loss observed for the Vanquish Neo-Orbitrap Ascend configuration relative to Easy-nLC-QExactiveHF, consistent with the generation of sharper chromatographic peaks by Vanquish Neo that places greater demands on MS duty cycle and chromatographic sampling under compressed gradient conditions (Fig.2B-C). Missing values in DIA ranged from roughly 2-7% and were elevated for mixtures with a larger dynamic range (Mix1, Ctrl) relative to mixtures with smaller inter-species abundance differences (Mix2) (Fig.2D) with a consistent <5% median coefficient of variation (%CV) across all the compared methods (Fig.S2D-E).

In regard to quantitative accuracy, DIA and MS3 outperformed MS2 and MS3RTS methods in the higher fold change-range with median log_2_FC close to the expected value of −2 for bacterial proteome, whereas all methods were consistently inaccurate at the difference of log_2_FC +1 (for yeast proteome) with highest relative accuracy achieved in TMT-MS3 (Fig.2E-F). As expected, TMT with MS2 was the least accurate in all regards due to ratio compression^21^ (Fig.2F). The deviation in accuracy appears to be dependent on summed protein abundances (MaxLFQ for DIA and signal-to-noise (S/N) TMT), with consistent dispersion observed at lower values across all compared methods and gradient lengths (Fig.S3A-B). In the lower FC range, where log_2_FC ±0.415 was expected, all TMT methods with 12 fractions (MS2, MS3RTS, MS3RTScl) struggled with accuracy (Fig.2F). This observation agrees with previous studies that employed ten TMT fractions versus ten DIA runs^22^, suggesting that accuracy in TMT experiments might be dependent on first dimension fractionation. However, this phenomenon could also be instrument dependent as TMT-MS3RTS on Orbitrap Ascend has previously been shown to incur a slight deviation in accuracy^23^. DIA on the other hand performed consistently across varying gradient lengths irrespective of the instrument setup suggesting that DIA is capable of robust quantification even in log_2_FC ±0.415 range independent on elution complexity (Fig.2E), specifically with >6 average data points/peak (Fig.S2C) as reported previously^20^.

Next, we compared TMT-MS3RTS datasets processed with Proteome Discoverer 3.2 and FragPipe version 23.0 (Fig.S3C-E). A similar comparison has previously been conducted, differing only slightly in parameters and use of an additional Chimerys module^24,25^. Our findings are broadly consistent with the earlier report^25^, showing approximately 95% overlap in protein quantification between the two analysis tools (Fig.S3C), although the correlation of measured fold changes was below 90% (Fig.S3D). FragPipe-derived intensities exhibited reduced accuracy relative to Proteome Discoverer’s signal-to-noise–based and TMT-purity corrected quantification, particularly for differential abundance ≤two-fold change (log_2_FC ± 1) (Fig.S3E) suggesting that TMT purity correction might be essential to accurate quantitation. We further assessed whether TMT correction or the presence of empty channels in the Proteome Discoverer quantification workflow affected results. Although both corrected and uncorrected approaches showed high correlation, omission of TMT correction introduced a systematic slope shift (y = 0.91x + 0.0072) and decreased quantitative accuracy (Fig.S3F) whereas presence of empty channels in the analysis did not seem to affect the overall quantitative accuracy (Fig.S3G). Together, our results provide evidence for deeper proteome coverage and quantitation achieved with TMT based methods as compared to LFQ-DIA within our set-up with similar quantitative accuracy achieved in LFQ-DIA and TMT-MS3. Moreover, the accuracy for MS3 methods may be dependent on TMT correction and the number of fractions, with less fractions and un-corrected TMT providing less quantitative accuracy. Overall, our findings support the use of TMT-MS3 (with or without RTS) for achieving deep and accurate proteome coverage on tribrid instruments, whereas DIA-based analyses may be better suited for hybrid systems where TMT reporter ions cannot be resolved in MSⁿ or corrected post hoc. The DIA approach, however, offers considerable speed and cost benefits when maximum depth and data completeness is not a primary requirement.

### Evaluation of dynamic proteomic screening with pSILAC-TMT and - DIA

To assess whether our optimized digestion from 96-well cell culture and acquisition strategies are compatible with dynamic proteome analyses, we designed a pulsed stable isotope labelling (pSILAC) experiment using two SILAC and threeSILAC labelling schemes to monitor protein synthesis. Cells were treated with thapsigargin, and its effects measured on protein translation (2SILAC) as well as recovery post-treatment (3SILAC) as compared to DMSO-treated controls (Fig.3A). Thapsigargin is an inhibitor of sarco-endoplasmic reticulum Ca(2+)-ATPase SERCA, induces unfolded protein response (UPR) and causes translation attenuation^26^.

**Fig 3:**
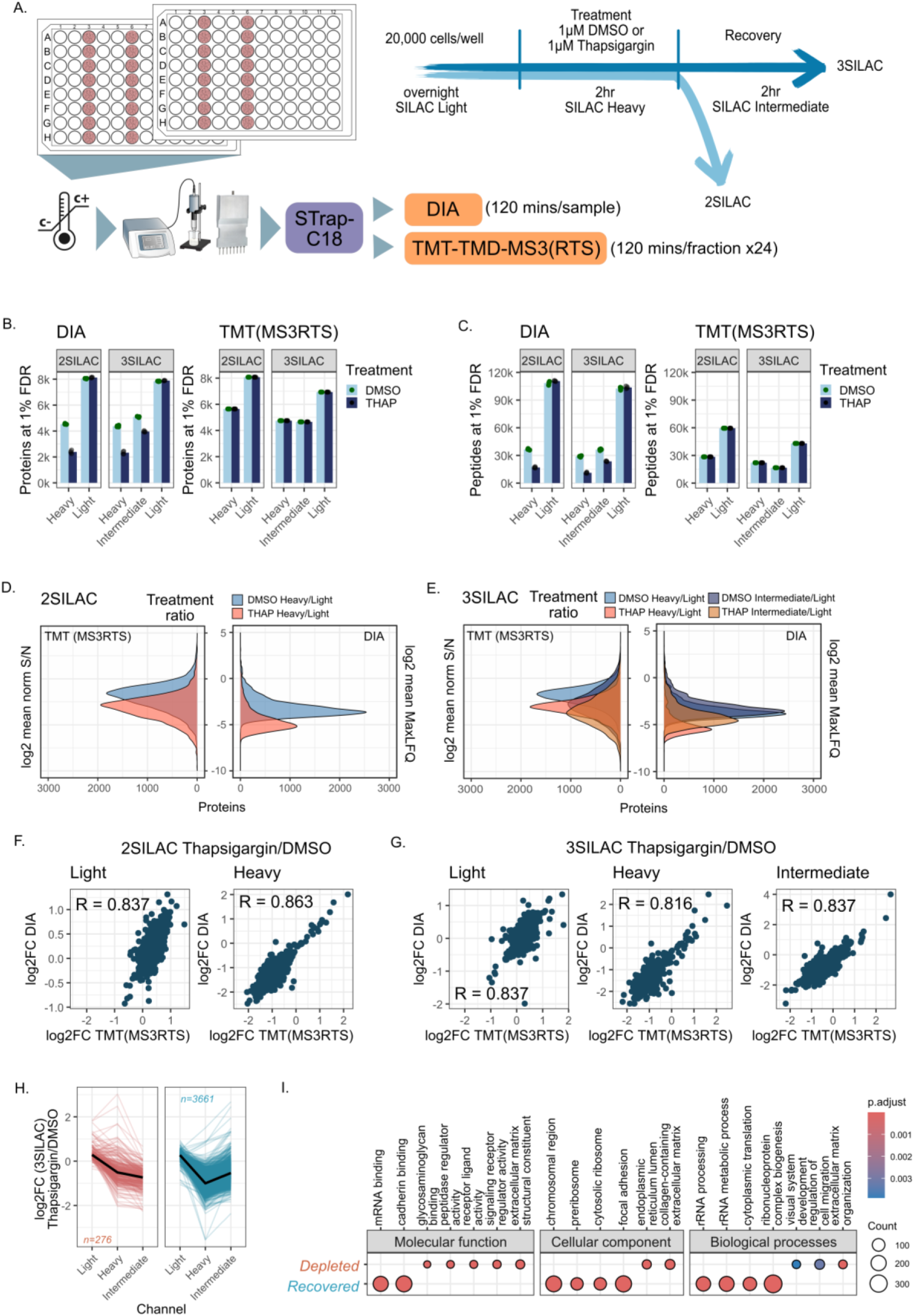
Evaluation of dynamic proteomic screening with pSILAC-TMT and -DIA. A. Experimental scheme for dynamic proteomics with pSILAC coupled treatment with DMSO or Thapsigargin (2SILAC) and recovery post treatment (3SILAC). 20,000 cells/well were grown overnight, treated for 2hrs and recovered for 2 hrs. The samples were processed with established STrap protocol and then either run as a TMTpro experiment (n=8 per condition, 24 fractions x 120mins) with targeted mass difference node activated (TMD) and real time search assisted MS3 scan or run as DIA experiment (n=4 per condition x 120mins). B. Number of Proteins and C. Peptides quantified across both DMSO (dark blue) and Thapsigarging (THAP) (light blue) treated cells in 2SILAC and 3SILAC setup across all included SILAC channels. D. Log2 distribution of mean ratio of Heavy/Light Protein abundance (S/N for TMT(MS3RTS) and MaxLFQ for DIA) for 2SILAC and E. Heavy/Light and Intermediate/Light Protein abundance for 3SILAC for DMSO and Thapsigarging (THAP) treated cells. F. Correlation of estimated log2 fold change (Thapsigargin/DMSO) for light and heavy proteome pool for 2SILAC and G. light, heavy and intermediate for (3SILAC) between DIA and TMT(MS3RTS) method. H. Log2 fold change (Thapsigargin/DMSO) of proteins quantified with all 3 isotopes in 3SILAC experiment, showing basal levels (light), treatment induced translatome (heavy), and post treatment recovery of proteins (intermediate). Left panel (brown) show proteins that remain depleted after removal of Thapsigargin and Right (turqoise) show proteins that recover after abolishing the Thapsigargin treatment by changing growth media. I. Gene ontology annotation of depleted and recovered pool of proteins showing their molecular function, cellular localisation and biological processes association from 3SILAC (TMT-MS3RTS) experiment.

The cells were processed with the previously established STrap protocol (this study, Fig.S4A) and the performance of the different measurement approaches compared by analysis using DIA on Q-Exactive HF and Lumos, or TMT-based MS3 acquisition with targeted mass difference (TMD)-enabled isotope selection^27^, with or without RTS, on the Ascend and Lumos respectively (120 min × 24 fractions; Fig.3A). Comparison of DIA and TMT-MS3RTS revealed distinct performance characteristics across SILAC plexes. DIA yielded consistent protein and peptide identifications across both 2SILAC and 3SILAC experiments, whereas TMT-MS3RTS exhibited a reduction in identification depth when moving from 2SILAC to 3SILAC (Fig.3B-C). Of importance, while the proteins quantified with DIA in the light channel expectedly remained unchanged across both treatments, a significantly lower number of proteins and peptides were quantified in thapsigargin treated samples, reflecting the profound effect on protein synthesis within cells, and consequently leading to missing values for quantitative statistics in DIA measurements. Conversely, TMT-based methods consistently identified all isotopes for both treatments with 2SILAC and 3SILAC setup, albeit with reduced log_2_ fold changes (Fig.S4B), making TMT-MS3RTS more amenable to quantitative differential abundance analysis without the need for imputation. This is also reflected in the ratio of heavy and intermediate SILAC labelled protein abundances relative to their light pre-treatment counterpart, wherein TMT ratios show more symmetrical Gaussian distribution across the dynamic range, and DIA MaxLFQ ratios show a sharp truncation of peaks on the lower end for both DMSO and Thapsigargin (Fig.3D-E). Moreover, the relative %CV for TMT-MS3RTS was significantly better for 2SILAC and marginally better for 3SILAC in comparison to DIA (Fig.S4C). For TMT-MS3 and DIA on the Lumos (Fig.S4D-E) the trend was similar to that observed with DIA on the Q-Eactive HF and TMT(MS3RTS) on the Ascend, however, the number of proteins and peptides are slightly lower for the same acquisition time as compared to TMT-MS3RTS on the Ascend or DIA on the Q-exactive HF. This agrees with previous in-depth comparison of the Lumos and the Q-Exactive HF-X, wherein Orbitrap MS^n^ acquisition on Lumos underperformed^28^.

More than 80% correlation was observed in the differential protein abundance profiles between thapsigargin- and DMSO-treated cells when comparing DIA and TMT(MS3RTS) analyses (Fig.3F-G) across all isotopes. In line with previous observations, proteins labeled with heavy isotope were consistently down-regulated following thapsigargin treatment^29–31^, whereas majority of intermediate isotope labeled proteins (3,661/4,656) showed recovery towards basal levels during the post-treatment phase (Fig.3H). These were mainly involved in translation/ribosomal machinery and RNA processing (Fig.3I). A small proportion of proteins (n=276) remained depleted even after removal of thapsigargin and were found to be mainly localized in extracellular matrix, secretory vesicles and endoplasmic reticulum lumen (Fig.3H-I).

Furthermore, we adapted the search settings for SILAC-TMT on FragPipe 23.0 with a custom roll-up of proteins from PSM.txt files and found a strong agreement amongst proteins identified and quantified with Proteome Discoverer 3.2 (Fig.S4F-G). Interestingly, while most proteins across all three isotopes were quantified with both search engines, FragPipe seemed to have a bias for higher heavy and intermediate identifications whereas Proteome Discoverer had a bias for light isotope labelled proteins (Fig.S4F). The isotope error correction feature within MSFragger-FragPipe 23.0, which is absent in Proteome Discoverer 3.2, is likely the cause of this discrepancy. Together, our results suggest a deeper coverage of dynamic proteomic screen incorporating 2SILAC with RTS-enabled MS3, whereas DIA achieves consistent coverage independent of the number SILAC channels.

## Discussion

In this study, we evaluate an accessible framework for deep and dynamic proteomic screening from 96-well cell culture formats using non-automated workflows and Orbitrap-based instrumentation operating below 50 Hz scan rates. By systematically benchmarking sample preparation strategies, acquisition modes, and data analysis approaches, we demonstrate that robust proteome depth and quantitative performance can be achieved without reliance on robotic automation or ultra-fast mass analyzers. These findings address a practical gap between state-of-the-art proteomics capabilities and the infrastructural realities of many laboratories. Plate-based cell culture assays are routinely applied within various contexts in cell biology and functional genomics. Yet, the limited protein yield per well often necessitates robotic liquid handling or a positive pressure platform to ensure robust and reproducible sample preparation. Here, we showed that comparable throughput and reproducibility could be achieved using standard laboratory equipment, such as a multi-channel pipette, 8-horn sonication adapter, deep well plates and a common plate-centrifuge. Among the evaluated workflows, both STrap with C18 clean up and Simplit2 combined with SDB-RPS enabled reliable processing of 96-well cell culture samples without automation, with latter being a more economical alternative while the former being the most robust for label-free and isobaric tagging based quantitative proteomics (Fig.1A-F, Fig.S1A-I). Although the Simplit2 workflow yielded lower proteome depth, increased inter-sample variability, and a higher proportion of missed cleavages relative to STrap, several factors could plausibly mitigate these limitations. Reducing the concentration of sodium deoxycholate in the lysate, employing higher-efficiency proteolytic enzyme, or transferring lysates from cell-culture plates to vessels optimized for proteomics could each improve digestion efficiency and reduce losses due to surface absorption. As an alternative low-cost strategy, n-dodecyl-β-maltoside (DDM), a mass spectrometry-compatible detergent previously applied in single cell proteomics^32,33^, may offer improved solubilization with reduced interference. In addition, low-cost alternatives for STrap using DNA purification Miniprep filters have been reported to achieve relatively better^34^, although their suitability for 96-well plate or higher throughput applications remain to be systematically evaluated. Of critical importance here are the plates used for cell-culture as there are various matrix options available for aiding cell adherence, which might however contribute to background contaminants or adsorptive losses. In this study, we therefore employed standard tissue-culture-treated polystyrene plates to minimize such confounding effects.

In our benchmarking with three-species peptide mixtures, DIA delivered consistently accurate quantification on both LC–MS platforms across active gradient from 120 to 45 min (12 to 32 samples per day; Fig.S3A, Fig.2E), although identification depth declined progressively as throughput increased. The sharper loss of identifications observed for the Vanquish Neo-Orbitrap Ascend configuration at the shortest gradients, which could be due to three factors i.e.: (i). lower automatic gain control (AGC) settings or shorter injection times for Ascend DIA scans can reduce the number of ions sampled per scan and thereby sensitivity; (ii) operating the Ascend at reduced MS2 resolving power to preserve cycle time sacrifices fragment separation and may lower identifications in crowded spectra; and (iii) narrower chromatography peaks on Vanquish Neo compared to Easy-LC at the same flow rate of 400nL/min. While we focused on maintaining comparable average numbers of data points per chromatographic peak across platforms, performance of the Vanquish Neo-Orbitrap Ascend configuration at shorter gradients could likely be further improved by optimizing the Vanquish Neo flowrate, Ascend DIA scan AGC and resolution. Irrespective of instrument tuning, we observed that missingness in DIA is strongly driven by sample dynamic range (Fig.2D, Fig.3B-C). Consequently, both experimental design and downstream data processing should explicitly account for this behavior, for example, through appropriate sample loading, conservative false discovery rate thresholds, sparsity-aware statistical frameworks, judicious use of imputation, or analysis platforms that explicitly model DIA missingness, such as Spectronaut^35^.

TMT workflows using MS3 (TMT-MS3, TMT-MS3RTS and TMT-MS3RTS with close out) delivered the deepest proteome coverage with precise quantitation in our evaluation and achieved near-complete data matrices. We did not see a substantial increase in proteins quantified with RTS closeout methods (2.45%), suggesting that the detection threshold likely plateaued with 12 fractions. However, close out method might be advantageous when using fewer than 12 fractions or shorter gradients for TMT multiplexed samples as it selectively limits MS3 triggering, thereby reducing redundant MS3 sampling of highly abundant proteins and reallocating duty cycle to lower abundance species relative to RTS method without close out (Fig.2C). These advantages of TMT can be further leveraged by incorporating higher-plex reagents (e.g., 32-plex)^36^, which allow more samples per run for a given acquisition time. A key practical consideration for TMT-based workflows is the cost of reagents, fractionation infrastructure and associated bench time imperative to reconcile their advantages. Additionally, proteome depth in TMT experiments may be compromised with increasing pSILAC plexing in dynamic proteomics, likely reflecting stochastic sampling effects arising from the expanded peptide isotope pool. By contrast, DIA remains largely agnostic to increase in isotopes if the fragment ions are effectively resolved (Fig.3B-C, Fig.S4D). A potential concern in DIA-based SILAC experiments is that isotopic pairs may be split across adjacent isolation windows, and potentially cause distortion in SILAC ratio due to differential peak sampling, as noted previously^37^. We reasoned that this potential distortion would not affect turnover measurements in pSILAC experiments as isotopes can be treated as separate protein pools and relied on channel-specific confidence metrics (Channel Q.value) implemented in DIANN v2.2 to separate the precursor isotopes prior to protein roll-up. Consistent with prior benchmarking, such an approach has been shown to preserve detection sensitivity and quantitative accuracy across large isotopic imbalances, including ratios down to 1:100 isotopic difference^38^, supporting the applicability of DIA to dynamic SILAC analyses. On tribrid instruments, ion trap can be utilised for accelerated DIA acquisition with narrower isolation windows and resonant CID-type fragmentation to achieve more data points per peak and deeper coverage^39^. However, resonant CID in ion trap is prone to lower mass accuracy relative to Orbitrap HCD^40^, and HCD fragmentation generates a richer population of singly charged y-ions^41^, particularly informative for resolving SILAC isotopes, especially when co-isolated and co-fragmented within the same DIA windows; thus, we favored Orbitrap-based HCD acquisition for isotope-resolved DIA in this study. Although DIA achieved slightly lower proteome depth than fractionated TMT workflows under the current instrumental setup, it required substantially less sample preparation time and incurred lower consumable cost. TMT reporter-ion quantitation can also be performed at MS2 level on hybrid Orbitrap instruments (for example, the Q-Exactive HF), which in principle may increase throughput and deliver comparable proteome coverage with fewer fractions or runs. However, as shown in Fig.2F, MS2-level TMT quantitation inherently affected by reporter-ion interference arising from co-isolated peptides, leading to substantial ratio distortion that typically requires post-hoc correction via computational approaches or experimental strategies such as baseline channel incorporation^21,31,42,43^.

For TMT data analysis, we used Proteome Discoverer and compared results to the freely available FragPipe (academic license). Both pipelines produced highly overlapping protein lists, but we observed small systematic differences in quantitative accuracy that we attribute primarily to TMT purity correction and differing handling of isotope errors. In our pSILAC–TMT data, FragPipe’s isotope-error correction improved identification and quantification of heavy/intermediate channels relative to Proteome Discoverer, suggesting that it is advantageous for dynamic-pSILAC with TMT. It will be worthwhile to test other freely available analysis tools that support TMT- correction (for example Comet^44^ and MaxQuant^45^), especially for SILAC-TMT with a ground truth dataset to more fully assess performance for dynamic-TMT designs. For DIA, we relied on DIA-NN 2.2, especially for its channel-aware scoring for plexDIA/pSILAC measurements and its improved FDR estimation compared to previous versions. Nevertheless, users should consider entrapment-based FDR estimation (as described previously^46^) especially in older DIANN versions and/or filter the report.parquet file for Q-value, PG.Q-value, Lib.Q-value, Lib.PG.Q-value and channel.Q-value (for SILAC) all set to < 0.01 along with Quantity.Quality > 0.8. A detailed guide on processing different SILAC experiments by Frankfenfield et al^47^ and the approach from this manuscript can serve as a primer for handling pSILAC DIA data analyzed with DIANN.

In conclusion, we present an accessible and cost-effective framework for deep and dynamic proteomic screening in 96-well cell culture plate, based on non-automated low-cost sample preparation workflows, widely deployed hybrid and tribrid Orbitrap instruments, and open-access analysis tools. By systematically comparing isobaric TMT- and DIA-based quantitation strategies for both global and pSILAC proteomes, we demonstrated the temporal resolution that can be achieved in understanding cellular trafficking and biological processes in a proteomic screen format. This framework can be further adapted for chemo- and geno-proteomics to achieve biologically important insights on older generation of instruments and still compete meaningfully in the race of ultra-fasts.

## Acknowledgement

The authors would like to thank Centre for Functional Proteomics (ZFP) Frankfurt and Qing Yu (UMass Chan Medical School), Vadim Demichev (Charité - Universitätsmedizin Berlin) and Fengchao Yu and Alexey Nesvizhskii (University of Michigan) for their valuable inputs on this manuscript. This work was supported by the Deutsche Forschungsgemeinschaft (German Research Foundation, DFG) for funding the LC-MS system (Easy nLC 1200, QExactive HF - Project-ID: 259130777, SFB1177 – Selective Autophagy and easy nLC1200, Orbitrap Fusion LUMOS - FuGG Project-ID: 403765277), SFB1531 Stroma-vascular damage control - project ID 456687919, SFB TRR387 Functionalizing the Ubiquitin System against Cancer – project ID 514894665, the European Fonds for regional development (REACT-EU, IWB-EFRE-program Hessen, 20008763) for the Vanquish Neo, Orbitrap Ascend LC-MS systems used in this study, the German Federal Ministry of Education and Research (BMBF) within the Clusters4Future PROXIDRUGS program (FKZ03ZU2109FC), the Hessian Ministry of Science and Research, Arts and Culture LOEWE Spitzenprofessorship, the Volkswagenstiftung, the Alfons and Gertrud Kassel-Stiftung, the Dr. Rolf M. Schwiete-Stiftung, and the Johanna Quandt-Universitäts-Stiftung. YL received a fellowship from Chinese Scholarship Council (CSC).

## Data availability

The .Raw files and processed files from Proteome Discoverer, FragPipe and DIANN can be found on ProteomeXchange under identifiers PXD073367 and PXD073468. Search results and R codes for data processing and visualization can be found on Zenodo DOI: 10.5281/zenodo.18925014.

