## Supplementary figures for "Evaluation of deep and dynamic proteomic screening strategies at sub-50Hz scan rate and without automation"

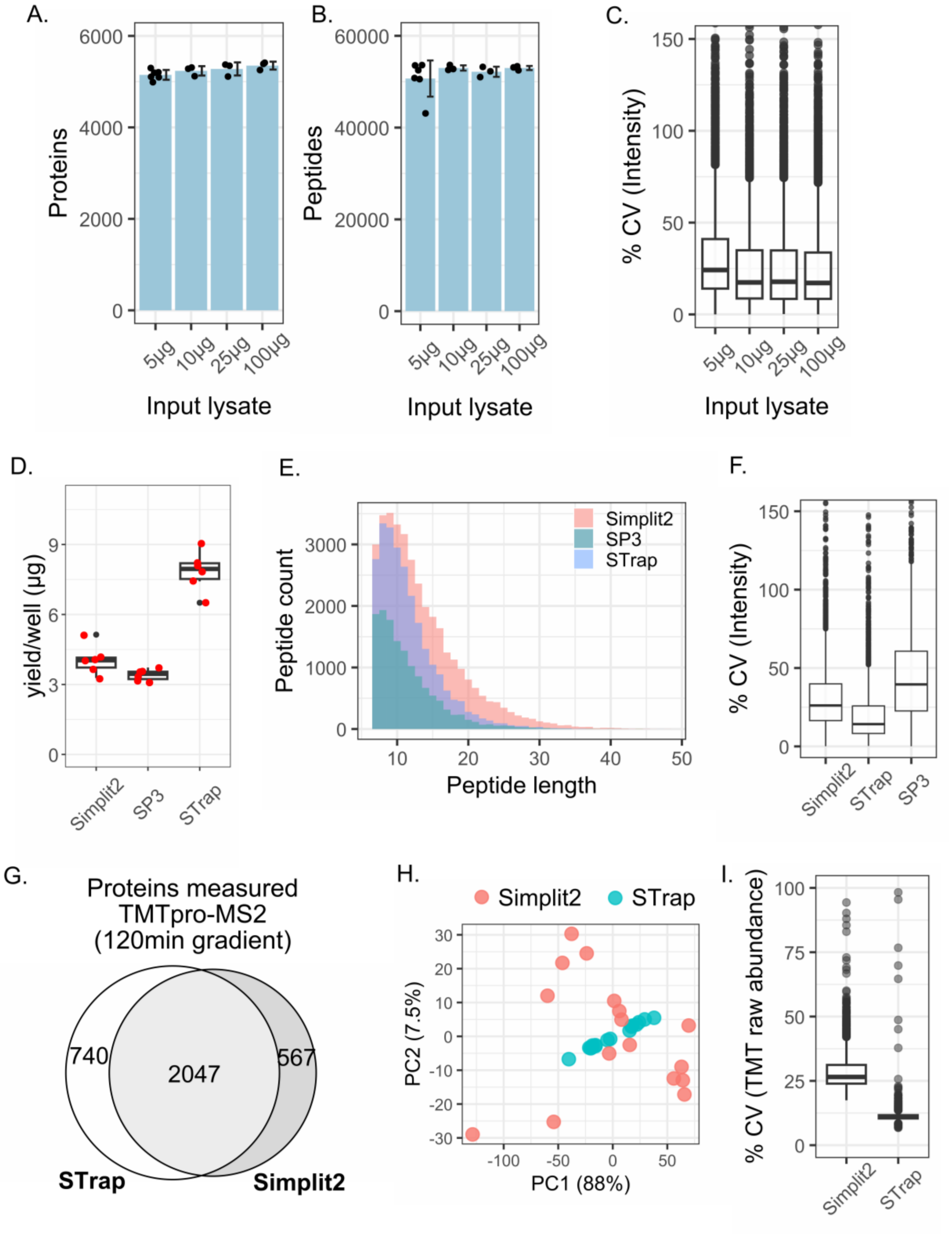


*Fig S1: Optimisation of high-throughput, non-automated sample preparation methods and TMT multiplexing of peptides for cells grown in 96-well plate.*

*A. Number of Protein and* B. *Peptides identified for HeLa lysate input amount 5µg (n=6), 10µg, 25µg, 100µg (n=3) processed with STrap 96 plate.* C. *Percent coefficient of variation in summed intensity of proteins for each input lysate in STrap plate with DDA-LFQ.* D. *Yield of peptides based on microBCA assay from a single* *well of a 96-well plate, seeded with 20,000 HeLa cells, grown over night (16hrs) and processed with either Simplit2, SP3 or STrap and cleaned with SDB-RPS (Simplit2) or C18 (SP3,STrap), both in 96-well format.* E. *Distribution of peptide length from tryptic digests and* F. *Percent coefficient of variation in summed intensity of proteins following Simplit2, SP3 and STrap and peptide cleanup.* G. *Overlap of proteins identified 20,000 HeLa cells, grown over night (16hrs) in 96-well cell culture plate (n=16) processed with Simplit2+SDb-RPS or STrap+C18, labeled with TMTpro and measured with MS2 - 2 hrs gradient on Q-Exactive HF.* H. *Principal component analysis and* I. *Percent coefficient of variation between the 16 TMTpro samples for Simplit2 and STrap2.*

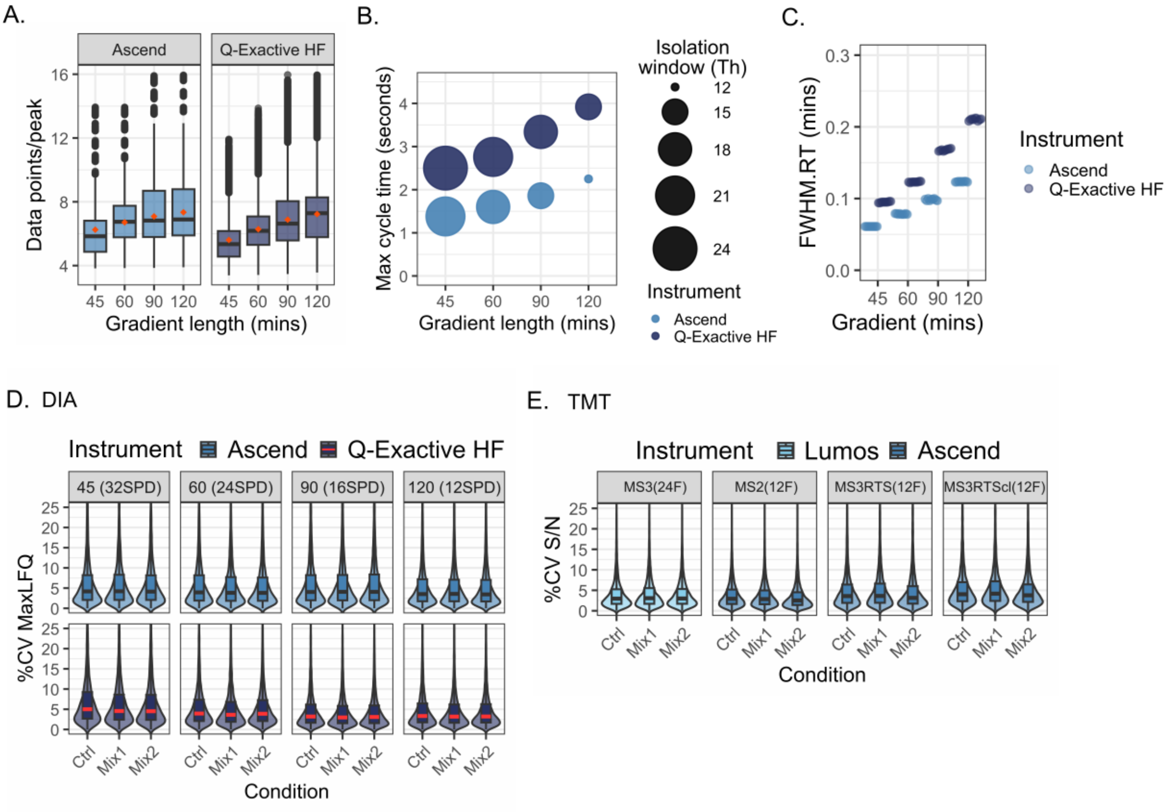


*Fig S2: Data acquisition parameters and quality statistics*

*A. Estimated datapoints/peak based calculated by taking the difference in retention time (RT.Stop and RT.Start) for each precursor multiplied by 60 and divided by the maximum cycle time for each method on Ascend and Q-Exactive HF. B. Max cycle time (seconds) for each gradient length and isolation window (Th) for DIA scan on Ascend and Q-Exactive HF. C. Estimated full-width half maxima retention time (FWHM.RT) per sample for each of the gradients run on Ascend and Q-Exactive HF as reported by DIANN 2.2.0. D. Percent coefficient of variation of DIA and E. TMT across all methods and instruments tested.*

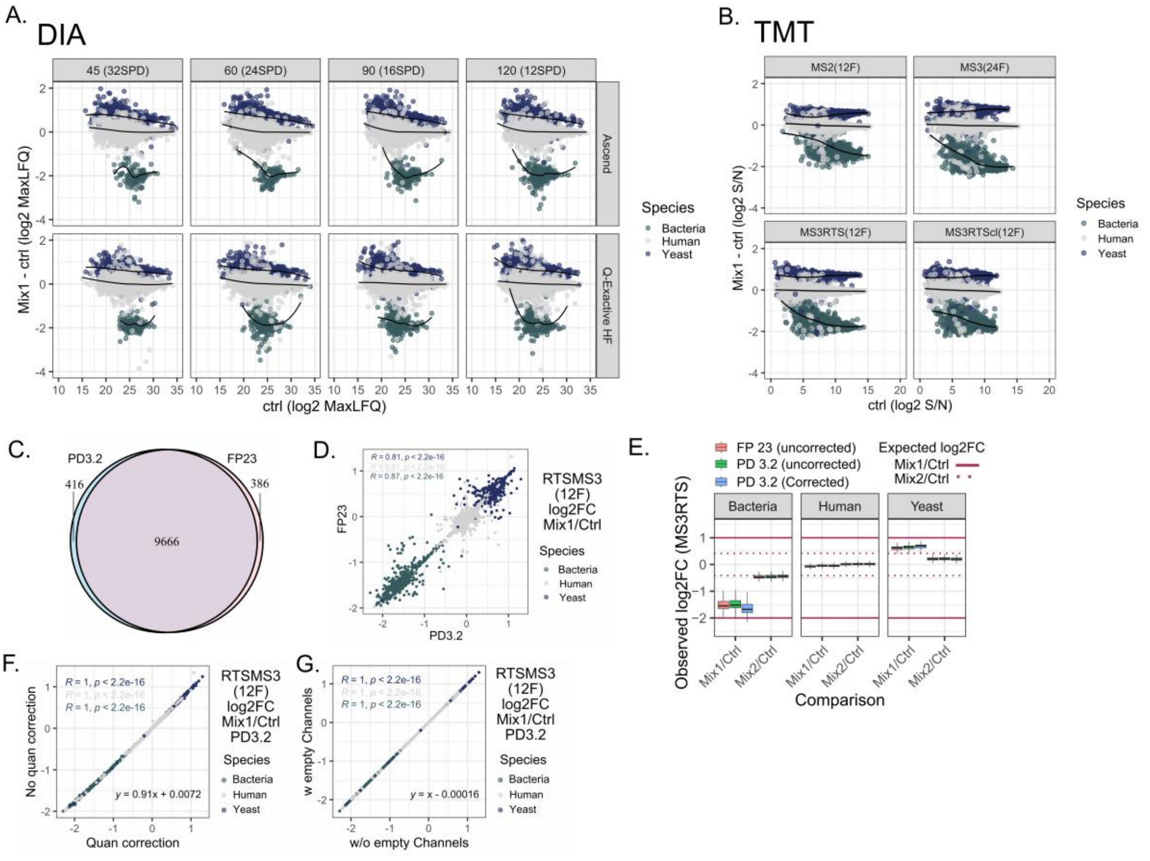


*Fig S3: Qualitative and quantitative statistics of DIA and TMT and comparison of TMT data analysis strategy.*

*A. Distribution of $\log_2$ mean MaxLFQ in Ctrl sample (x-axis) against the difference in Mix1 and Ctrl log2 values (y-axis) with varying amounts of yeast (blue) and bacteria (green) peptides mixed in equal human (grey) peptide background separated over 45, 60, 90 and 120 mins gradient length and analysed via DIA on Ascend and Q-Exactive HF. B. Similar to (A) but Proteome Discoverer derived signal-to-noise ratio (S/N) for TMT values for four TMT methods tested on Lumos and Ascend. C. Overlap of proteins quantified with Proteome Discoverer (PD) 3.2 and FragPipe (FP) 23.0 in TMT-RTSMS3 samples (12 fractions without peptides/protein close out. D. Correlation between $\log_2$ fold change in Mix1 as compared to Ctrl sample, based on TMT intensities derived from FragPipe and signal-to-noise ratios derived from Proteome Discoverer 3.2, dots separated by species as in (A) but only for RTSMS3 samples. E. Observed (boxplot) and expected (line) $\log_2$ fold change between Mix1/Ctrl (solid) and Mix2/Ctrl (dotted) for the 3 species based on intensity values from Fragpipe 23.0 and signal-to-noise from Proteome Discoverer 3.2 either corrected for TMT-Purity or uncorrected in consensus "Quant correction" for RTSMS3 samples. F. Correlation in Mix1/Ctrl $\log_2$ fold change in Proteome Discoverer 3.2 output with and without TMT-purity correction for RTSMS3 samples. G. Correlation in Mix1/Ctrl $\log_2$ fold change in Proteome Discoverer 3.2 TMT-purity corrected output with and without 2 empty channels in analysis quantitation method.*

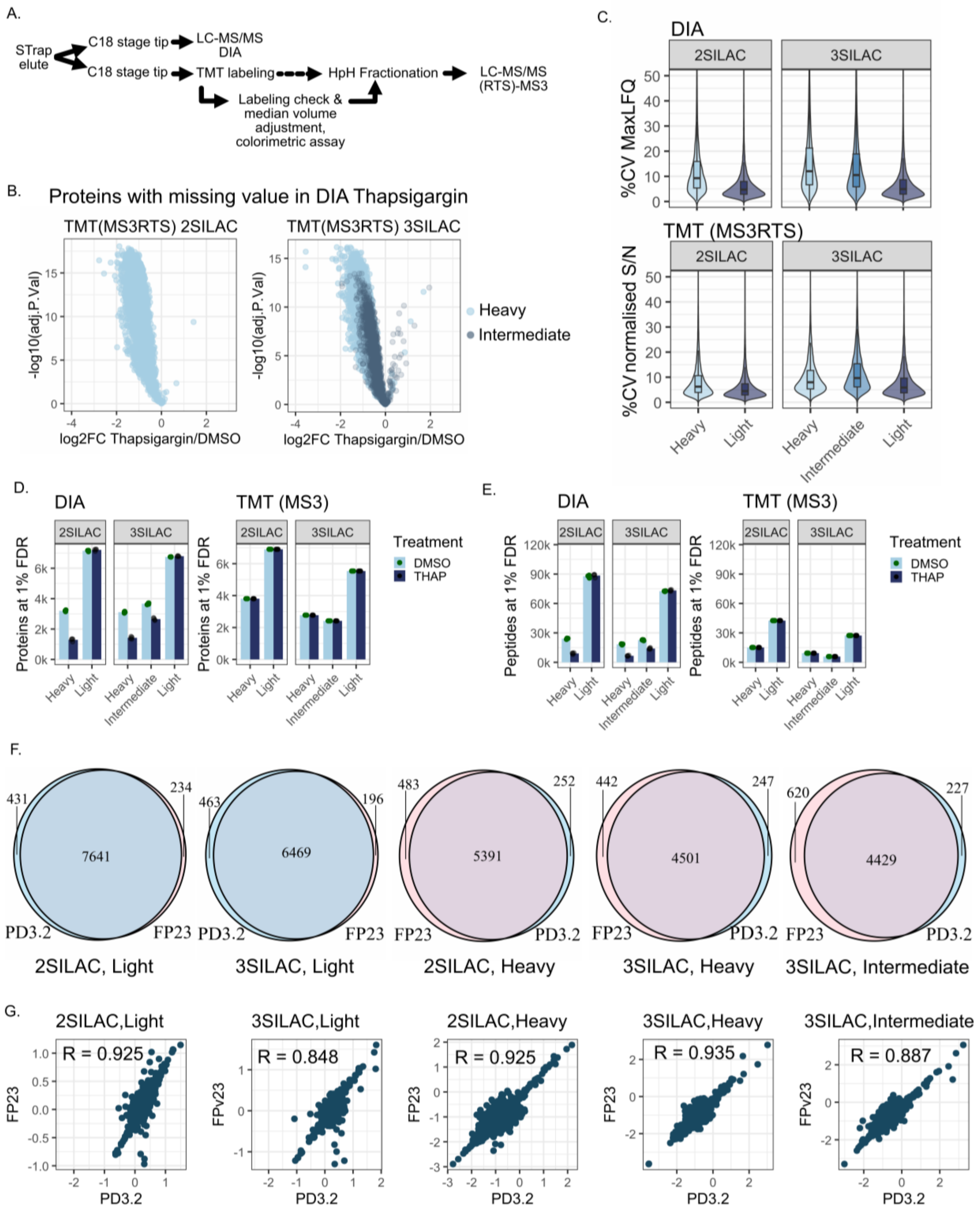


*Fig S4: Comparison of DIA and TMT methods for pSILAC.*

*A. Schematic workflow of steps involved from peptide elution from STrap to LC-MS measurement. For DIA STrap eluted peptides are dried, cleaned with C18 and ready for measurement. For TMT, additional steps such as TMT labeling, labeling check, volume adjustment, colorimetric assay and fractionation were performed before the samples were measured on MS. B. $Log_2$ fold change of proteins identified in heavy* *(for 2SILAC - left panel) or heavy and intermediate (for 3SILAC - right panel) isotopes in TMT(MS3RTS) method of Ascend but have missing values in Thasigargin sample when run as DIA on Q-Exactive HF. C. Percent coefficient of variation of MaxLFQ values for DIA (top) and signal-to-noise values for TMT(MS3RTS) (bottom) for all isotopes in 2SILAC and 3SILAC experiment. D. Number of proteins and E. Peptides quantified with the same samples as Fig.3 analysed on Obritrap Lumos in either DIA mode of SPS-MS3 mode for TMT labeled fractions. F. Overlap and G. Correlation of $log_2$ fold changes of proteins detected as either light, heavy or intermediate (based on psm assigned modification) in Proteome Discoverer 3.2 and FragPipe 23.0 in 2SILAC and 3SILAC TMT(MS3RTS) data.*

*Table.S1: Sheet 1: Data acquisition parameters, Sheet 2: Gradient parameters for DIA.*
